# Understanding artificial mouse-microbiome heterogeneity and six actionable themes to increase study power

**DOI:** 10.1101/778043

**Authors:** Abigail R Basson, Alexandria LaSalla, Gretchen Lam, Danielle Kulpins, Erika L Moen, Mark Sundrud, Jun Miyoshi, Sanja Ilic, Betty R Theriault, Fabio Cominelli, Alexander Rodriguez-Palacios

## Abstract

The negative effects of data clustering due to (intra-class/spatial) correlations are well-known in statistics to interfere with interpretation and study power. Therefore, it is unclear why housing many laboratory mice (≥4), instead of one-or-two per cage, with the improper use/reporting of clustered-data statistics, abound in the literature. Among other sources of ‘artificial’ confounding, including cyclical oscillations of the ‘cage microbiome’, we quantified the heterogeneity of modern husbandry practices/perceptions. The objective was to identify actionable themes to re-launch emerging protocols and intuitive statistical strategies to increase study power. Amenable for interventions, ‘cost-vs-science’ discordance was a major aspect explaining heterogeneity and the reluctance to change. Combined, four sources of information (scoping-reviews, professional-surveys, expert-opinion, and ‘implementability-score-statistics’) indicate that a six-actionable-theme framework could minimize ‘artificial’ heterogeneity. With a ‘Housing Density Cost Simulator’ in Excel and fully annotated statistical examples, this framework could reignite the use of ‘study power’ to monitor the success/reproducibility of mouse-microbiome studies.

## INTRODUCTION

Laboratory mice are critical to understanding human biology in a variety of fields, from inflammatory bowel diseases, neurology, and cancer, to microbiome and nutrition. In the current era of microbiome research, multiple factors are becoming evident as sources for confounding. Integrating microbiome science into animal research necessitates that experiments control for confounding derived from emerging artificial factors, especially the ‘cage microbiome’,^1-5^ which we recently discovered causes ‘cyclical microbiome bias’ due to the periodic accumulation of excrements in mouse cages.^1^

Understanding the factors that contribute to research heterogeneity will address this need. Primary factors causing artificial analytical heterogeneity and low study power include putting many mice into one cage, having insufficient cages per group, and using statistical methods that assume multiple mice in a cage are independent instead of clustered observations.

In statistics and science, heterogeneity is a concept that describes the uniformity and variability of an organism, a surface, or the distribution of data. Sources of study heterogeneity can be natural or artificial. Artificial heterogeneity refers to study variance introduced by humans or anthropological factors, including animal husbandry and the ‘cage microbiome’, which non-uniformly affect mouse biology. Fundamental to hypothesis testing, data heterogeneity determines which statistical methods are needed to decisively quantify if two independent naturally-heterogeneous groups, truly differ. To appropriately select statistics controlling for cage-clustered data, scientists must be aware of study details, namely, which data points belong to which mice and respective cages in a dataset or published figure. Unfortunately, these details are often omitted during analysis and in publications, and misconceptions on heterogeneity, husbandry and analysis may exist among leading research organizations.

To exemplify that scientists are under pressure and need recommendations to prevent bias and improve animal research quality and reproducibility, in the USA, the National Institutes of Health (NIH), a major federal funding institution, implemented a mandate in 2014 on ‘Rigor and Reproducibility’^6-8^ which assures research funding is constrained unless researchers prove that they consistently yield reproducible results. Our report seeks to illustrate concepts on study power and intra-class correlation among mice in a cage to support a framework based on six actionable themes to increase study reproducibility.

Concerning study power, two concepts of expected validity exist: internal and external validity. Both refer to the statistical expectation that results from a given study are true, reproducible, and not by random chance if a study is repeated locally (internal), or in another setting (external validity).^9-11^ Intrinsically, experiments have high internal validity if appropriate statistics and power are applied, and if data clusters and confounders are avoided. Studies with experiments in different settings (microbiota, mouse lines) are more likely replicable; but experimental reproducibility requires appropriate power. Validity thus depends on the study power, which is the probability of not making a type II error (fail to reject false null hypotheses in favor of true alternatives). Power is a statistical measure from 0 to 1, with 1 indicating highly-powered studies. While power 0.5 yields statistically haphazard results (‘tossing a coin’), powers >0.8 indicate optimal chance for replication. Power increases with large sample sizes (more mice), but decreases with clustering of animals in cages by introducing a ‘cage effect’, and intra-class correlation coefficient (ICC) complexity to the analysis of cage-clustered data. The negative impact of cage clustering is maximum when all mice of a study group are housed in one cage because it is impossible to differentiate ‘real’ from ‘confounding cage effects’. The negative impact of clustering is reduced when more cages, with *fewer mice per cage*, are used per group (*‘less mice-per-cage is more’*).

Despite the 5-year-old NIH mandate, the public and federal perception on mouse research reproducibility is often negative.^7,12^ However, to our knowledge, there are no scientific studies ***i)*** confirming that research reproducibility is an ongoing issue, ***ii)*** defining what role perceptions and academic husbandry practices play on reproducibility, or ***iii)*** predicting the implementability of potential solutions to increase study power, if proposed. To refine our understanding on research heterogeneity, study power and reproducibility, our study objectives were to ***i)*** verify research methods heterogeneity in current literature, ***ii)*** quantify current perceptions on mouse husbandry and microbiome using a survey, ***iii)*** identify potential areas of solution using a Delphi-based strategy, and ***iv)*** to quantify the potential implementability of an evidence-based framework of six Recommendation themes to cost-effectively increase study power using a grading scale based on perceived clarity, benefit and recommendability.

As an accompanying practical set of tools, we *<u>also</u>* created ***i)*** a simple housing density cost calculator in Excel that can be used by scientists to determine whether less animals per cage, or more cages per experimental group suit research budgets, and ***ii)*** and provide graphical examples and a fully annotated statistical code to compute and report analysis of cage-clustered data, and power, for both single- and clustered-caged mice. Post-hoc study power calculations were deemed cumbersome and non-informative in the past,^13^ but more sophisticated user-friendly software now provides emerging methods to compute such important statistics,^14^ which we propose should be used to infer and objectively monitor power and reproducibility across mouse research at large.

## RESULTS

### Husbandry heterogeneity and cage-cluster effects are pervasive in current literature

To identify husbandry factors capable of influencing gut microbiome and study reproducibility, especially mice per cage (MxCg) and mice per group (MxGr), we reviewed 172 recent studies selected from PubMed searching ‘diet-microbiome-mice’ (**Figure 1**). From 865 articles published over the past 10 years, 93% were published in the last five years (**Supplementary Materials;** https://figshare.com/s/9d0b963e287944233cb1). Of concern, most studies failed to report in sufficient detail aspects of animal husbandry (*e.g.*, cage density/sanitation frequency, diet sterility) making the study of cage-effects and confounding challenging to assess (**Figure 2, Supplementary Figure 1**). Although 57% of the studies originated in China and USA (n=52, 30%), it is remarkable that almost 60% of studies across all countries failed to report animal density (*i.e.*, MxCg). Of the 72 studies that reported density, 30% (22/72) have highly cage-clustered data; reporting experiments with 5 MxCg. Slightly encouraging, 18% of studies housed mice at lower densities of ≤2 MxCg, which is ideal because it increases study power by decreasing cage effects (**Figure 2A-C**). Although low animal density could be perceived as an expensive practice, density practices did not correlate with gross domestic product (GDP; yearly US$/capita) implying that national wealth is not a driving factor for housing mice individually during experiments. Irrespective of wealth, it was reassuring to identify scientists who publish studies stating that they exclusively housed mice individually in Belgium, Taiwan, Italy, Finland, Korea, France, Brazil and Japan^15-45^ (**Figure 2 D-E**).

**Figure 1.**
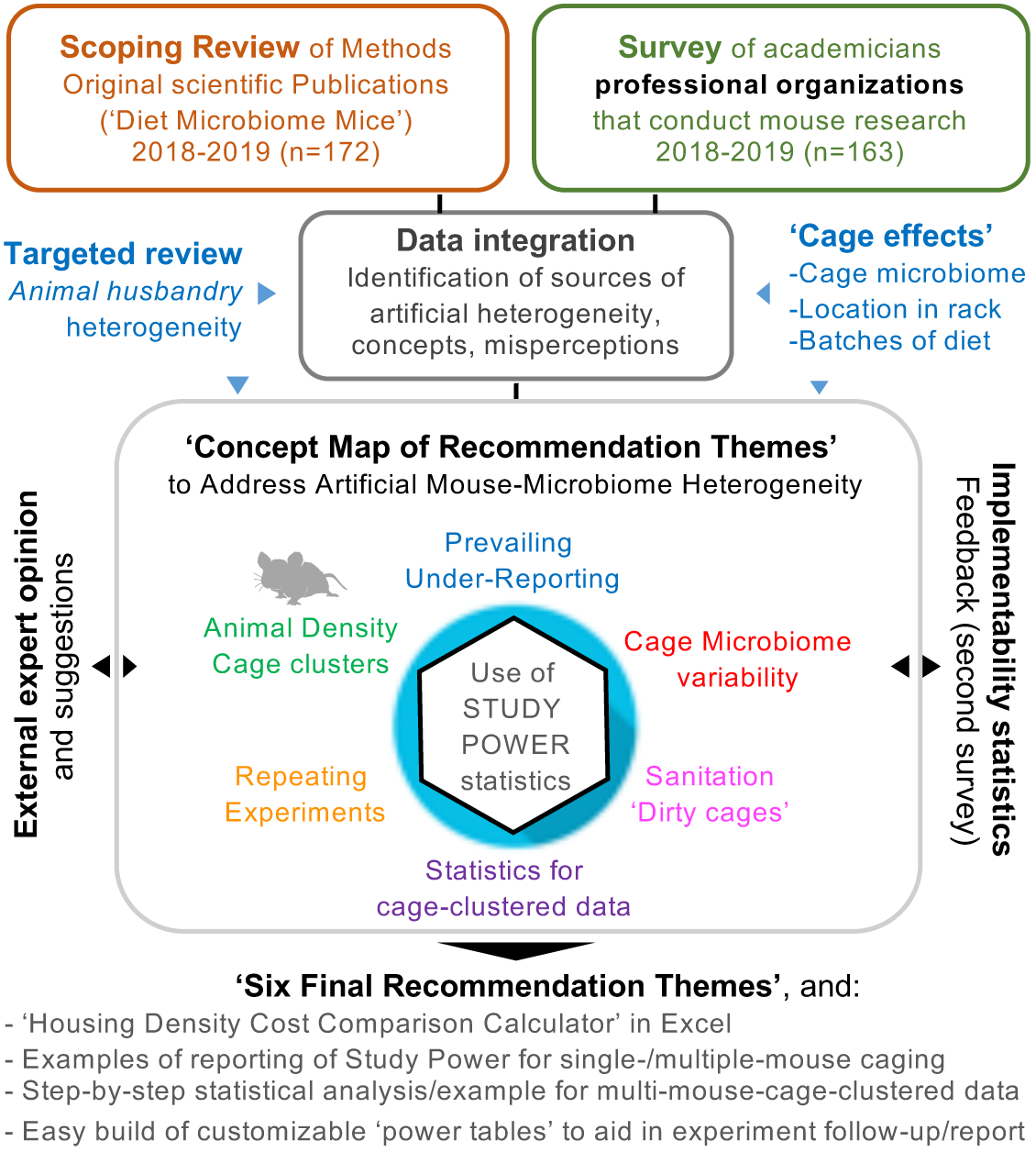
Study design to understand artificial heterogeneity in mouse microbiome research.

**Figure 2.**
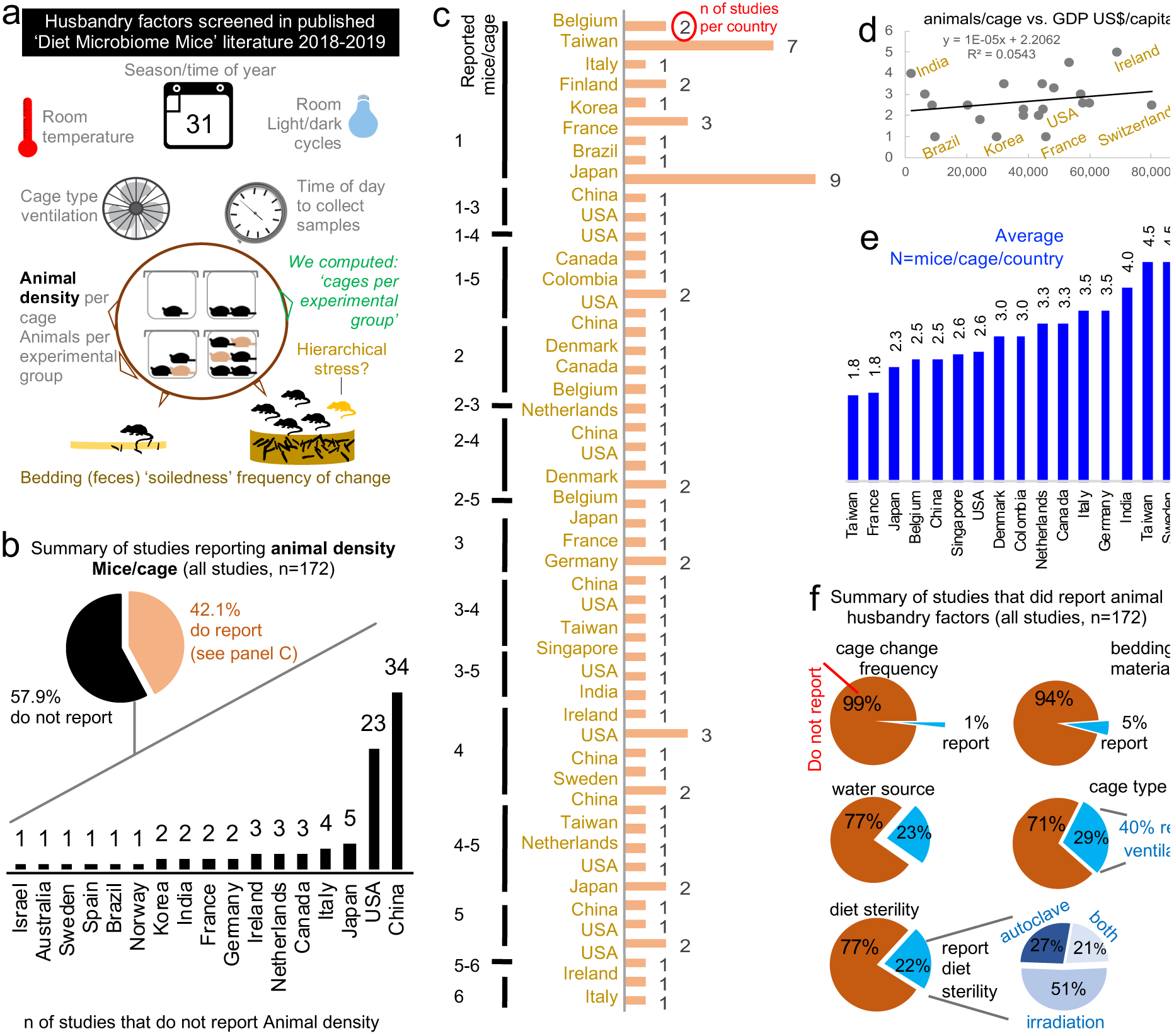
Literature on ‘diet, gut microbiome & mice’ illustrates ongoing animal density problematics. Published methodologies illustrate variability in husbandry and inconsistent animal density across studies as a major source of cluster-confounding. **a**) Schematic representation of factors screened from the methods and results section in peer-reviewed publications. **b**) Distribution of studies that did and did not report animal density. Pie chart shows that most studies (58%) do not report how many animals were housed per cage. **c**) Ranking shows number of studies per country based on the number of studies reporting animal density (78 of 172 reported). **d**) Correlation between number of MxCg and GDP US$/capita. Note that the country’s GDP does not correlate with number of MxCg suggesting experimental animal density practices are not related to wealth of a country. **e**) Average MxCg used in experiments represented by country. **f**) Summary of studies that reported cage change/sanitation frequency, bedding material and diet sterility (including method for diet sterilization; autoclaving, irradiation & dose used). Note that more studies reported ‘cage type’ (e.g., plastic flexible film, metal wired, Plexiglas, etc.) than those which reported ‘sterility of diet’ (25% vs 21%). Only one study reported ‘time of fecal collection’ (see complementary data in Supplementary Figure 1).

Several husbandry aspects contribute to cage-cage variations and cause cage effects (see **Supplementary Table 1-2**). Therefore, it is difficult to substantiate whether the significant effects identified in any given study, where all mice in a group were housed in one single cage (decreasing study power), were truthfully due to the experimental intervention and not from the random distribution of cage effects in a laboratory (**Figure 2F**). To quantify the potential for ‘cage effect confounding’, we used the ‘total number of cages per group’ (TCgxGr) as a quantitative estimate (see **Methods**) to determine the prevalence of studies that conducted experiments using only a few cages per group. Estimates indicate that studies used on average 4.4±3.2 TCgxGr (notice large SD), of which 39% (28/72) generated data derived from only 1-2 TCgxGr (**Supplementary Figure 1**).

Given that cage clusters decrease study power,^46-48^ experiments conducted with low animal density, ideally one MxCg, and the reporting of TCgxGr deserves to be highlighted as an exemplary habit. Despite available reporting guidelines,^49^ data illustrates that inadequate reporting of methodological details in published literature continues in 2019, diminishing the ability to replicate studies. To complement guidelines, we propose to consider using a standard verbatim paragraph-style format to unify reporting and facilitate future meta-analyses (see below **Recommendation Theme on ‘Reporting’**).

### Expertise differences across scientific organizations surveyed

To further advance our understanding of husbandry heterogeneity, we applied an online survey to academicians (**Supplementary Figure 2**). After contacting over 2000 professionals, a total of 166 participants started the online survey. One-hundred and sixty-three (97%) surveys were completed and used for analysis. The majority of respondents were from USA (133; 81%, 95%CI=74.3, 87.6) and participants reflected individuals with leading roles in science (Assistant Professors, Professors, Veterinarians) within the DDRCC, AALAS and GNOTOBIOTIC organizations (see **Methods**). The GNOTOBIOTIC respondent set had a smaller number of faculty/veterinary directors or managers (vs. Postdocs) compared to the DDRCC group (p=0.087, 61.4% vs. 78.8%, Odds ratio [OR] = 95% CI=0.82, 5.7) but included slightly more participants with access to germ-free (GF) animals compared to DDRCC (p=0.083, 95.5% vs. 84.6%, OR = 3.82, 95%CI=0.69, 38.5, **Figure 3A-B**). Multi-and single-cage GF isolators (used as a proxy for state-of-the-art equipment and knowledge) were most frequently used as a GF-caging system among those with GF facility access. Collectively, demographic analysis indicates that although statistically different, all groups had comparable levels of expertise, access to state-of-the-art facilities and knowledge (note p-values and wide 95%CIs; see **Figure 3C-D**) which is important to inferring that the perceptions acquired herein are relevant to current research.

**Figure 3.**
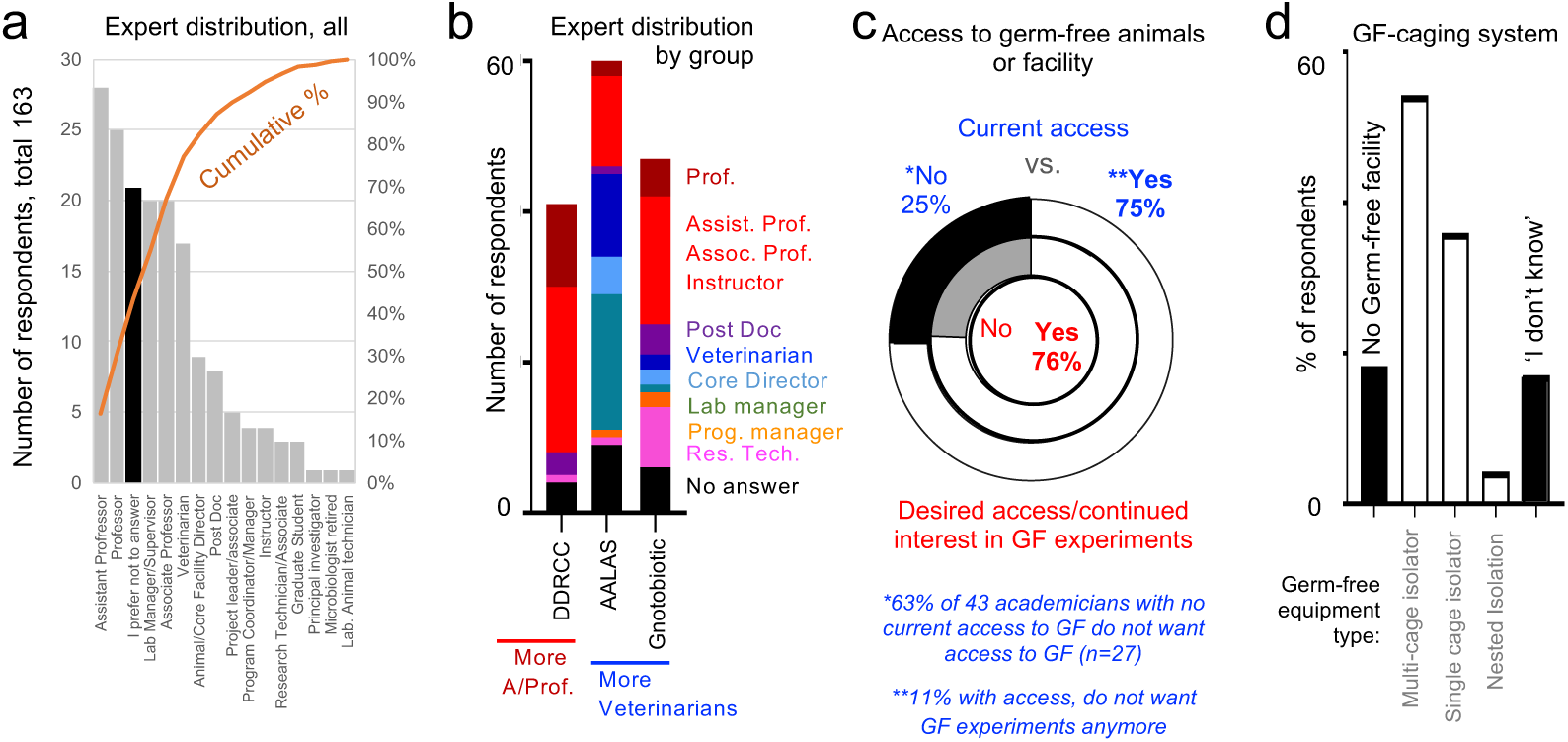
Demographics of surveyed professionals on ‘animal husbandry in microbiome research’. **a**) Pooled distribution of job descriptions categorized based on information provided by all respondents. **b**) Distribution of job descriptions by the three largest groups of participants. Notice that the DDRCC group has the largest proportion of faculty (from instructors-to-full professors) participating in the survey, but all groups were composed of academicians with comparable job descriptions. More veterinarians and project leaders were observed in the AALAS and Gnotobiotic listserv groups. **c**) Distribution of participants who reported having current access to GF animals or facilities (outer pie circle chart) and that would like to have access, or continue working with, GF animals/facilities (inner circle chart). Notice that the majority of participants are expected to have high levels of expertise and understanding of GF mouse facilities, husbandry, and microbiome knowledge. * and ** indicate subgroups who would like (or not) to change their current GF research trends. **d**) Distribution of respondents who did or did not know about the presence of GF facilities in their institution, and the types of caging system used. This question contextualizes the knowledge of respondents in terms of GF equipment/systems.

### Scientific organizations rank similarly 15 husbandry factors that affect the mouse microbiome

To determine whether differences in knowledge/practices or perceptions on animal husbandry exist due to the professional nature of each organization, we asked participants to rank, from 1 to 5 (least to most important), how important each of 15 husbandry factors contribute to variability in mouse research (“*Rank how important you believe each of the following 15 aspects contribute to microbiome research variability)*”. Using ‘*diet composition*’ as a positive control (as diet affects gut microbes), we found that all groups of professionals ranked each parameter similarly (mean of ranks for all participants across factors, Kruskal-Wallis p>0.05).

Except for *‘diet composition’*, ranked 1^st^ as ‘very important’ by the majority of respondents (>75%), there was marked heterogeneity in response patterns at the individual level (**Figure 4A**). Importantly, perceptions of individuals did not cluster within their professional affiliation, suggesting that the organizations surveyed ‘think’ alike. Instead, we identified ‘patterns of beliefs/perception’ in academia that reflect ‘types of individuals’, with a given set of research practices in mind (beliefs), that differs from their peers within their organization (**Figure 4A-C**). For example, although ‘*coprophagia’* ranked 4^th^ overall as a ‘very important’ factor to microbiome variability, fewer than 40% of participants ranked *‘number of animals per cage’* (ranked 8^th^) and *‘cage change frequency’* (ranked 9^th^) as aspects ‘very important’, even though coprophagia contributes to microbiome confounding depending on the extent of *‘cage bedding soiledness’* (ranked 12^th^), which depends on *‘number of animals per cage’* and *‘cage change frequency’*.

**Figure 4.**
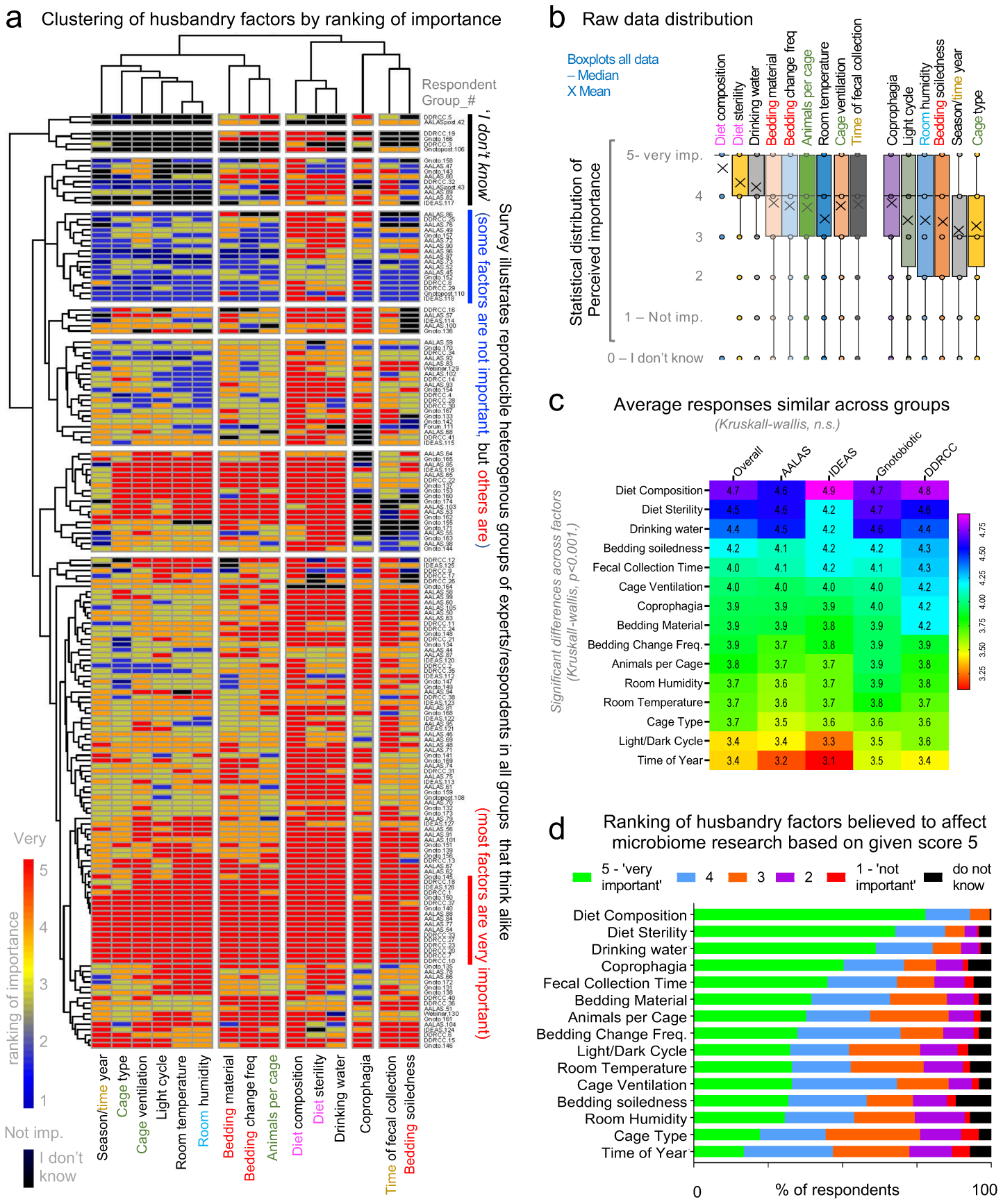
Ranking of 15 factors believed to cause microbiome research variability is reproducible. **a**) Heat map shows respondent perceptions on the importance of various animal husbandry factors in microbiome research variability. The heterogeneity across respondent perceptions illustrates that individual thinking is not related to institutional affiliation. **b**) Boxplots show raw data ranking distribution of respondent perceptions on the importance of various animal husbandry practices. **c**) Heat map shows the overall ranking of variables according to institution. **d**) Stacked bar graphs show overall ranking of variables. Note that diet composition, sterility and drinking water were identified by >50% of individuals as ‘very important’ contributors to microbiome research. Note the discordance between coprophagia (ranked 4^th^) to that of bedding soiledness (‘dirtiness’) and the importance of cage change frequency.

In the studies reviewed, aspects deemed ‘very important’ by survey respondents were not always reported, while ‘less important’ factors were frequently reported. This discordant pattern of thinking-reporting was further illustrated by individual perceptions on ‘*bedding type’* (*e.g.*, corncob vs. non-edible wood shavings), ‘*cage ventilation’* type, ‘*room temperature’* and *‘room humidity*’, all of which contribute to cyclical bedding microbial overgrowth (which selects for aerobic microbes in cage bedding) and thus cage-cage microbiome variability.^1,2,48^ Beliefs agreement was identified between ‘*diet composition’ ‘diet sterility’* and *‘water source’* (top 3 ranked factors) illustrating that dietary intake is perceived as a collective of all aspects consumed orally, including the microbial content of diet (**Figure 4D**). Most respondents do not think *‘cage type’* (ranked 14^th^) is important. The majority of reviewed studies (**Figure 2F**), however, reported cage type in their methods, while the ‘very important’ aspect of *‘diet sterility’* was described in only 22% of studies reviewed. Of concern, the *‘time of year/season’* was the least important aspect believed to influence the microbiome (ranked 15^th^); however, we have shown that cross-sectional metagenome experiments conducted in separate seasons produce contrasting results when assessing the role of *Helicobacter* spp. in spontaneous Crohn’s disease-like ileitis in mice,^3^ implying that repeating experiments across seasons may yield unreproducible results over time.

As a recommendation, repeating experiments to build composite datasets, which often occurs across seasons, should be conducted with caution unless we understand the effect of season on the microbiome and animal physiology (see **Recommendation Theme on ‘Repeating Experiments’**).

### Diet-dwelling microbes and homogenizing cage microbiome variability before experiments

With sub-sterilizing radiation protocols, diets have variable microbial composition even within the same batch.^1,2,50^ Survey questions interrogated basic knowledge relevant to irradiation and the degree of diet sterility. When asked whether standard irradiated commercial diets for mice were sterile, 67% answered that such diets were *‘sterile’*. Although diet sterility depends on the irradiation dose, in the case of commercial diets, companies employ a single, standard dose, insufficient to achieve GF-grade sterility. Of note, no studies reviewed reported irradiation dose when reporting *diet sterility*. Thus, unless certified as sterile, diets used during mice rearing and experiments expectedly contain potentially confounding microbes, primarily spore-formers and gamma-radiation resistant bacteria and fungi.^51^ The random distribution of diet-dwelling microbes, bedding-dependent microbial overgrowth and other cage effect factors are sources of microbiome divergence^52^ and bias that accumulate across cages as animals are reared and aged before, or during experimentation.

Since there is no consensus on one single approach to control for cage-cage microbiome variability before using mice in experiments, we surveyed which methods are used by scientists.^52-55^ Despite evidence that co-housed mice have varying microbiome patterns^56,57^ and the recent evidence of cyclical bedding-dependent bias,^1^ the most popular combination of methods used to control for cage microbiome variability was *‘cohousing’*, ‘*use of mixed bedding’* and *‘increasing the number of animals per cage’* (**Figure 5A**). The least frequently used method was *‘fecal homogenization’* (animals exposed to a composite of feces harvested from all mice), yet this method is arguably the simplest and most effective in homogenizing cage microbiome variability (see **Recommendation Themes on ‘Cage-cage microbiome variability BEFORE experiments’** and **‘Dirty cages and time-of-sampling DURING experiments’**).

**Figure 5.**
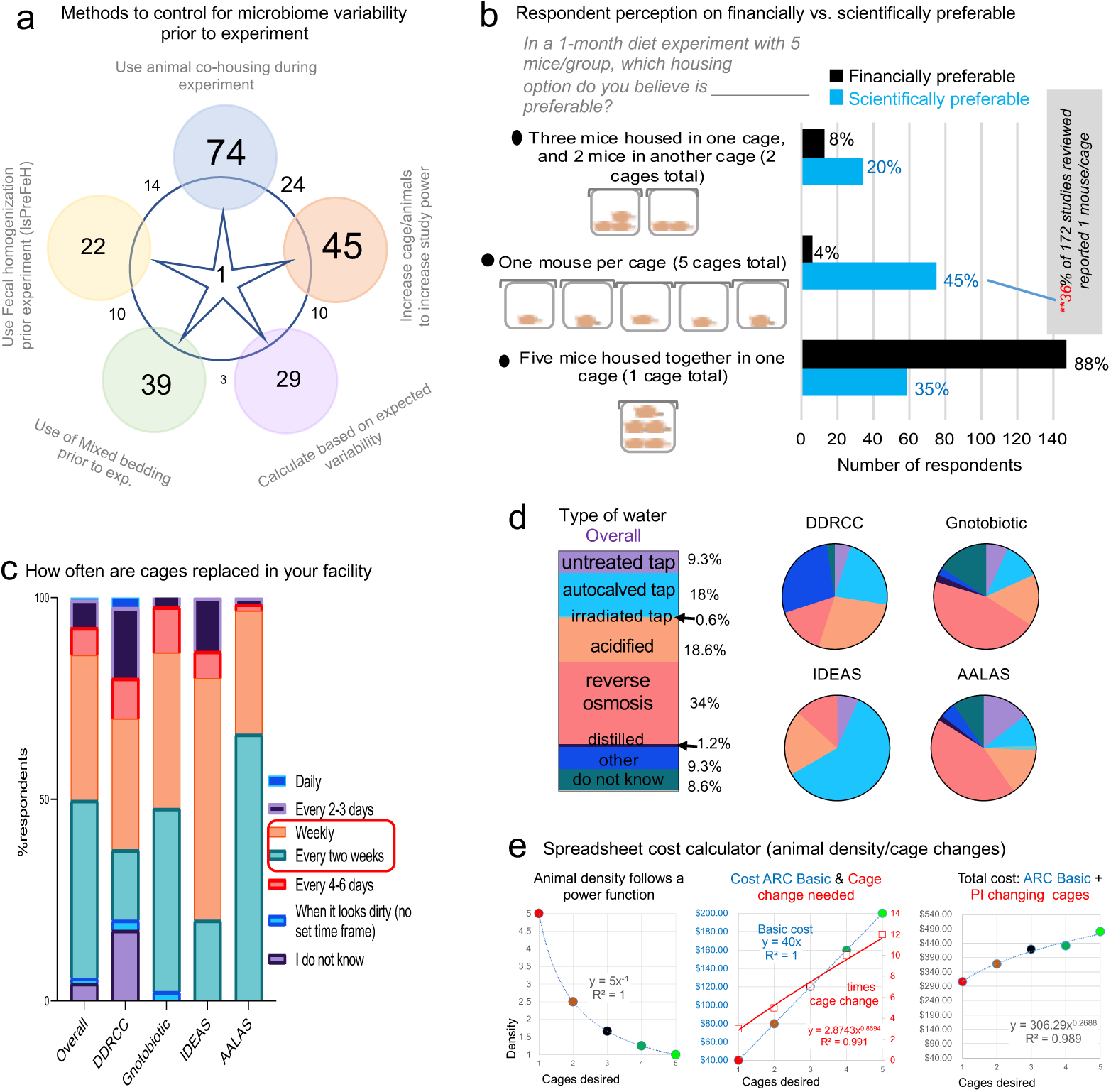
Survey responses for animal husbandry practices and cost. **a**) Venn diagram (n of respondents) on ‘popularity’ of various methods used to control cage-cage microbiome variability prior to the experiment. Note ‘fecal homogenization protocol’ compared to others. **b**) Perception contrast between the ‘financial’ and the ‘scientific’ preference when asked what animal density was preferable for a 1-month dietary experiment. Of interest, 88% and 35% of the survey respondents believe that 5 MxCg is financially and scientifically preferable than housing fewer animals per cage. **c**) Stacked bar plots show ‘cage change frequency’. Most facilities change cages weekly or every 2 weeks. **d**) ‘Water-type’ in facilities (8.6% ‘did not know’). Note wide array of water sources, including untreated tap, autoclaved tap, acidified tap and reverse osmosis, all of which affect the gut microbiota.^79^ **e**) Cost analysis example using a customizable spreadsheet calculator (**Supplementary File 1**). Notice the power function correlation between ‘number of cages desired’ in a study and ‘animal density’ with the linear costs of husbandry due to payment of ‘basic costs’ in an Animal Resource Center (ARC) and the presumed costs of cage handling by a technician paid by the Principal Investigator (PI).

### Clusters and scientific-financial discordance when housing five mice in a study of five mice

To interrogate whether cost is a contributing factor to animal housing density practices, we posed two identical multiple-choice questions that differed only by the assumption of financial vs. scientific preference. The first question asked, “*In a 1-month diet experiment with 5 mice/group, which housing option do you believe is FINANCIALLY preferable?* while the second question replaced the capitalized word ‘*FINANCIALLY’* with *‘SCIENTIFICALLY’.* The three possible answers were, using ‘5 cages’, ‘2 cages’, or ‘1 cage’. The majority of participants believe it is both scientifically (54%) and financially (95.7%) preferable to maintain cages with higher animal density (2-3 or 5 MxCg), which, of concern, introduces cage cluster effects.^58^ Thus, studies with 5 mice are underpowered as they consist of only 1-2 cages; commonly seen in studies reviewed. Intriguingly, while 45% (95%CI=37.3, 52.6) of respondents think that it is more scientifically appropriate to have 1 MxCg, the same individuals do not think that this practice is economically feasible (**Figure 5B**), which reflects current literature where only 15% (95%CI=9.6, 20.3) of studies reported exclusively housing 1 MxCg (see **Figure 2C**).

Considering that the majority of respondents’ facilities implement weekly or every 2 weeks ‘*cage change’* protocols, with a wide array of drinking water sources across facilities (**Figure 5C-D**), our data suggests that cage change/sanitation (via ‘cage microbiome’), and animal density could contribute greatly to artificial heterogeneity in mouse research.

To address concerns of cost regarding the number of MxCg in context to ‘*cage change frequency’*, we developed an Excel spreadsheet ‘Housing Density Cost Comparison Calculator’. Graphical cost-effectiveness analysis illustrates that a higher number of MxCg requires more frequent cage changes (**Figure 5E**, available as https://figshare.com/s/377fa429bd8cc405fc1b). Overall, costs increase when comparing 5 vs. 1 MxCg linearly over a continuum of cage cluster possibilities, therefore conducting highly clustered underpowered studies is not necessarily cheaper. When considering response patterns regarding financial vs. scientific feasibility of animal housing density, we show that the heterogeneity in respondents’ perceptions is not attributed to institution but instead to professional organization (**Figure 6A-F**).

**Figure 6.**
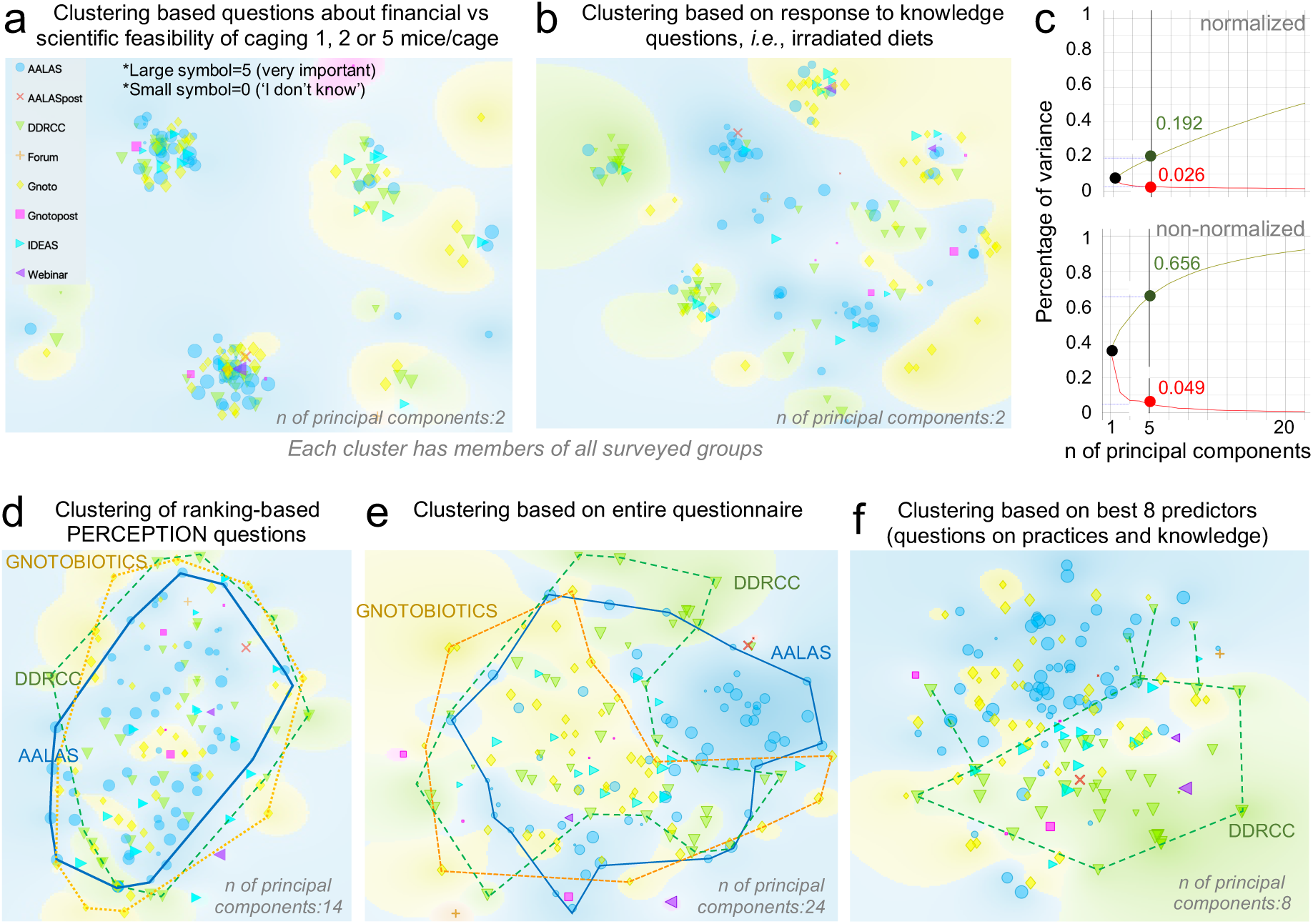
Beliefs on ‘husbandry and microbiome research variability’ are similar, but professional organizations differ in response to questions on practices and knowledge. Normalized principal component analysis of survey respondent data. Superscript asterisks: large or small symbols depict the individual response of each participant when asked how important ‘animal density’ was as a factor in influencing the gut microbiome.**a**) Clustering-based questions about financial vs. scientific feasibility of caging 1, 2 or 5 MxCg. Notice that each cluster (type of response patterns) contain individuals from all professional groups, i.e., AALAS. **b**) Clustering-based knowledge questions, i.e., irradiated diets. Notice the same pattern as in panel A, suggesting that response heterogeneity is not due to group. **c**) Normalized and non-normalized percentage of variance in entire data set explained by the maximum number of components (questions; n=24) using “animal density” as outcome for prediction (which cannot be achieved as large and small symbols occur throughout plot). d) Cloud representation of collective influence of the 15 questions to predict group separation. **e**) Clustering based on 15 ranking-based PERCEPTION questions + 11 Knowledge, Financial vs. Scientific feasibility, access to facilities and practices. Although clusters of individuals collectively think very similarly and slightly different than the rest, analysis indicate that the different clustering for certain areas in the plot is due to differences in answers related to ‘type of facilities’, or practices that are more common among certain groups of professionals. **f**) Best achievable clustering of individuals based on relief F scores to predict animal density shows surveys from different groups are distinct.

Although scientists could argue that statistical methods exist to control for clustering,^58^ our analysis of literature indicates that scientists do not implement cluster-statistics. Since cluster-statistics are not trivial to implement (*e.g*., R Statistical Package ‘clusterPower’^59^), we provide technical guidelines on how to account for unbalanced MxCg designs, ICC and low sample size using clustered-data statistics (see **Recommendation Themes 5-6 on ‘Animal density, clusters, ICC, and power’**).

### Implementability of a multi-theme framework to favor study power and reproducibility

To objectively determine if the ‘Recommendations’ described below (supporting a multi-theme actionable framework, **Figures 1** and **7A**) were ***i)*** clearly drafted as a sentence (*sentence clarity*), ***ii)*** had the potential benefit to improve power and reproducibility (*potential benefit*), and ***iii)*** were deemed appropriate for readers to recommend to others (*would you recommend it?*), we asked active academicians and scientists conducting research to grade each recommendation and provide comments to create an ‘implementability grade metric’ (**Supplementary Table 3**). To quantify whether the obtained implementability grades were significantly different from random responses, we compared the distribution of grades to that of a random generator of 30 numbers, from 1-10.

**Figure 7.**
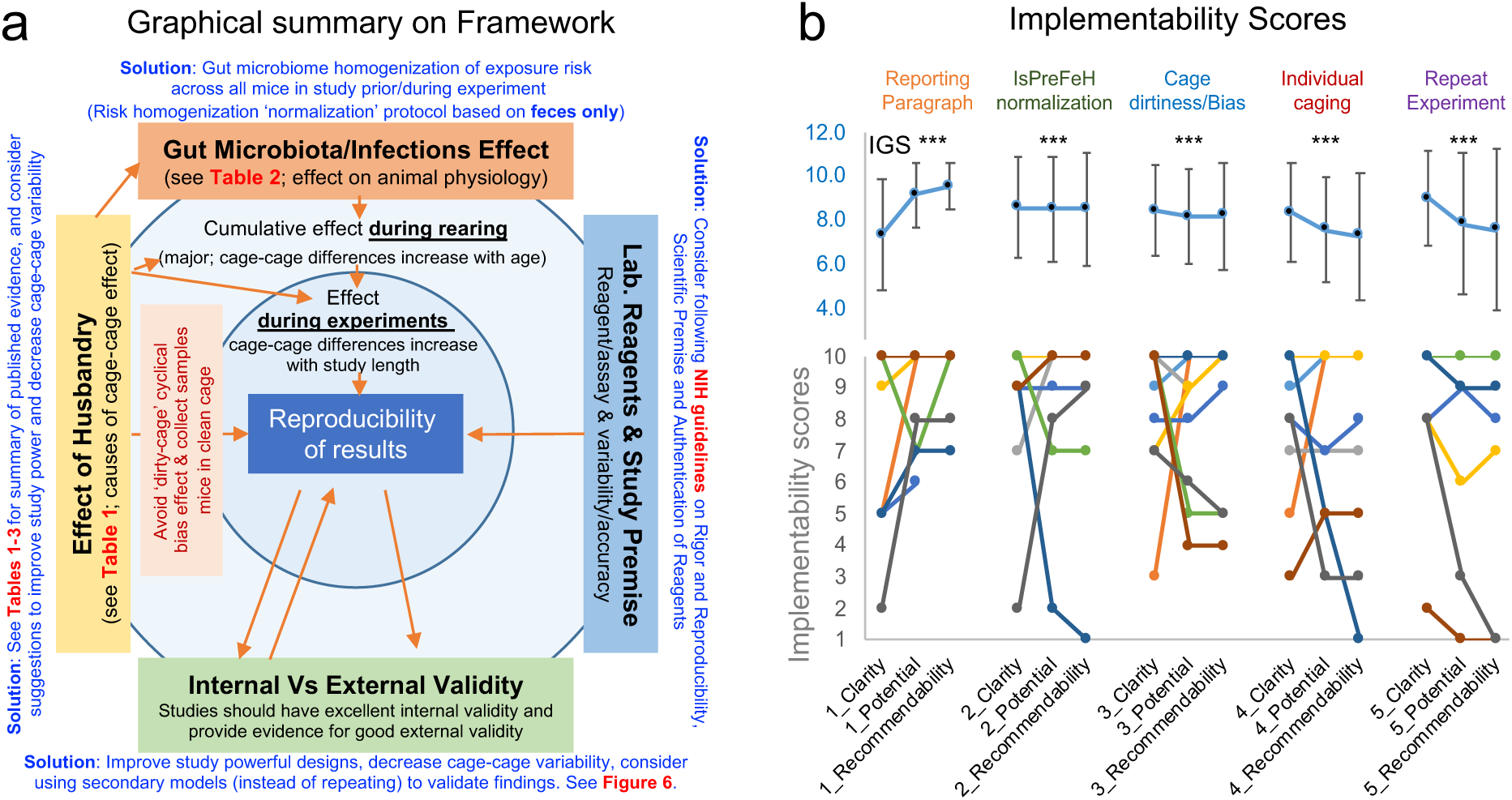
Implementability of recommendation theme framework. **a**) Framework integrating NIH guidelines, and our recommendations. **b**) Implementability grades scores (IGS) for each recommendation. Asterisks indicate IGS were statistically higher than random simulations (see statistics details in-text). Line plot connects individual grades. Notice that people who disagree with sentence clarity tend to disagree interpreting the potential benefits. High mean grades indicate high potential for implementability.

A grade of 1 indicates poor, while a grade 10 means outstanding. Of great practical value for the multi-theme framework proposed, analysis indicated that, collectively, all recommendations are very likely to be implemented by scientists (mean grade, 8.02±1.4 vs. random grade 5.0, n=20, t-test p<0.001; **Figure 7B**).

The wording of the final recommendations, underlined with ‘*<u>quotation marks and italics</u>*’ reflect the improved version of the expert-graded sentences and comments received during the grading phase. See all comments in **Supplementary Table 4**, and a synthesis of the peer-reviewed studies supporting the framework in **Supplementary Table 5**. The implementability statistics, rationale (extended version in **Supplementary Table 6**), and goals for each Recommendation are described below.

### Recommendation theme 1 on ‘Reporting of diet and husbandry factors’

Reproducibility will occur only if critical study details are provided in published literature. Our review of studies combined with the high number of ARRIVE guideline^49^ citations (>3000) indicates that while ‘checklists’ may improve reporting quality, they do not ensure reporting with sufficient/consistent detail. A template paragraph for reporting would enforce uniform transparency, reproducibility, and enable rapid data mining for future meta-analyses, widely used to help guide the practice of medicine, but scarcely used in basic science. We **recommend** the ‘*<u>Use of a paragraph-style template to report detailed diet and husbandry factors consistently and reproducibly (e.g., macronutrient, diet sterility), publishable as accompanying ‘Supplementary Materials</u>”* (see reporting template in **Supplementary Table 7**). The **goal** is to minimize reporting with insufficient detail or details that are open to interpretation, yet still suffice standard reporting checklists/guidelines^49^. The expert-prediction for **implementation** is significantly high (grade, 8.7±1.2 with 99.5% probability of being significantly higher than random in 96.7% [n=29] of t-test analysis conducted for 30 simulations with 30 random numbers, mean t-test p=0.005±0.012). Note that ‘text-recycling’ is currently allowed (when clearly justified) based on current code of ethics in scientific publishing.^60-62^

### Recommendation theme 2 on ‘Cage-cage microbiome variability BEFORE mouse experiments’

Fecal bacterial profiles can differ widely between cages within a single mouse strain housed under identical conditions and occurs even across mice produced for experimentation in contained breeding colonies.^2,3,63^ Our survey demonstrated that although scientists implement strategies to control for cage microbiome variability before experiments,^2,55,56^ there is ample variability of arguably reproducible method combinations used across organizations. A fecal homogenization protocol wherein all mice are administered a composite of freshly collected feces via oral gavage for 3 days,^54^ has been shown to effectively minimize inter-cage gut microbiota heterogeneity before experimentation.^54,55,64^ We **recommend** the ‘*<u>Use of a fecal matter-based microbiome normalization protocol (e.g., by orally administering a homogenous pool of feces from a group of mice intended for experimentation to all the mice at baseline prior to starting the study) to homogenize the microbial exposure risk across all mice intended for an experiment, and thus reduce the cage-cage microbiome variability that naturally occurs as animals age during intensive production of animals for research and experiments.</u>’* The **goal** is to normalize microbiome variability that accumulates across cages over the lifespan of mice before experiments. The expert-prediction for **implementation** is significantly high (grade, 8.5±0.04; 98.25% probability of higher score vs. random; significant in 86.6% of simulations, t-test p=0.018±0.03). Described in 2014 as ‘Inter-subject Pre-experimental Fecal Microbiota homogenization’ (IsPreFeH),^54^ this revised microbiome ‘normalization’ protocol, which excludes use of soiled bedding material, in combination with a reproducible protocol for oral gavage of microorganisms,^65^ is a scalable solution.

### Recommendation theme 3 on ‘“Dirty cages” and time of sampling DURING experiments’

Our survey showed ample heterogeneity in timing of mouse cage sanitation protocols despite recent studies indicating that bedding soiledness (‘dirtiness’) contributes to periodic variations in gut microbiome via contact/coprophagia.^1,52^ Mouse experiments would benefit if conducted with cages having reduced animal density (1-2 MxCg) with biological samples systematically collected from clean cages at the same time of day to avoid diurnal variation.^66-68^ We **recommend** to ‘*<u>Prevent the uncontrolled accumulation of animal excrements in the cage, **i)** house a homogeneous number of animals per cage (ideally at low density, 1 mouse/cage), **ii)** adjust frequency of cage sanitation based on animal density, and **iii)** collect samples 1-2 days after mice have been in clean bedding/cages, because coprophagia and ‘dirty cages’ affect the mouse physiology and microbiota.</u>’* The **goal** is to minimize the uncontrolled permanent contact of mice with their (decomposing) feces. The expert-prediction for **implementation** is significantly high (grade, 8.3±0.15; 98.7% of probability of higher vs. random; significant in 96.6% of simulations, t-test p=0.014±0.024). Given that coprophagia (not relevant to humans) and excrements in cages may cause bedding-dependent cyclical microbiome bias,^1^ frequent cage replacements (increases with animal density,^1^ **Supplementary Figure 3**), studying/sampling mice in clean cages and/or the use of slatted floors^69^ deserve emphasis.

### Recommendation theme 4 on ‘Repeating experiments in different seasons’

As reflected by the literature reviewed and the misconceptions documented in our survey, little is known about the effect of time of year/season on mouse research heterogeneity.^3,63,70^ Since it is almost impossible to control for seasonal variation within long-term, or multiple short-term experiments spanning over several seasons, it is important to take measures to improve measures taken to improve study variation/reproducibility over time (*e.g.*, food batch, inter-experiment IsPreFeH). We **recommend** to ‘*<u>Plan and execute statistically powerful designs and do not repeat underpowered (cage clustered, low sample size) experiments in different seasons (because several unforeseen factors affecting animal husbandry are challenging to detect and control for in diet and personnel)</u>.’* The **goal** is to control for the variable effect of season on study reporting and heterogeneity using well-powered designs. The expert-prediction for **implementation** is significantly high (grade, 8.1±0.76; 96% probability of higher score vs. random; significant in 76.6% of simulations, t-test p=0.04±0.062). We acknowledge that at times replication is desirable, and also that ‘poor breeding colonies’ often yield insufficient mice to perform final experiments. In this context, it is advisable to store fresh-frozen feces anaerobically (−80°C with/without cryoprotectants; 7%-DMSO, 10%-glycerol) from initial experimental mice for the colonization of newly available mice, and to store sufficient vacuum-packed diet (−20°C) and supplies to last across experiments.

### Recommendation theme 5 on ‘Animal density, clusters, and study power’

First, our scoping review identified numerous laboratories publishing clustered MxCg data with few cages/groups, without the verification of study power/sample sizes, or use of statistics for clustered-data. Then, our survey and cost simulator showed financial-scientific discordance among scientists when deciding animal densities. Unless higher densities are scientifically (not only financially) justifiable, housing 1 MxCg could yield more costeffective and powerful study designs by increasing the number of cages and minimizing the need to use advanced statistics.^47,71^ We **recommend** to ‘*<u>House one mouse per cage (unless more mice per cage is scientifically justifiable) and increase the number of cages per group (instead of few cages co-housing many mice which results in cage clustered-correlated data, lower study power and requires more mice to compensate for study power loss) to maximize the experimental and statistical value of each animal as a test subject during experimentation</u>.’* The expert-prediction for **implementation** is moderately significant (grade, 7.7±0.56; 91.4% probability of higher score vs. random; significant in 63.3% of simulations, p=0.086±0.13). The **goal** is to maximize the scientific/test value of each mouse by promoting individual housing, emphasizing that social stress has been equally demonstrated, irrespective of sex, for single- and socially-housed mice,^72,73^ and to promote the use of study power through cost-effective, reproducible experiments. As expected, this recommendation elicited the most heterogeneous responses, reflecting a partial reluctance to modify current animal density practices (**Figure 7B**). To promote implementation and facilitate the accuracy/reproducibility of reports, we provide three graphical examples of why/how-to compute and report power/sample sizes for any completed experiment using single-caged mice and intuitive open-access software (‘G*power’^74^ in **Figure 8A-C**, R^75^, and our STATA code below).

**Figure 8.**
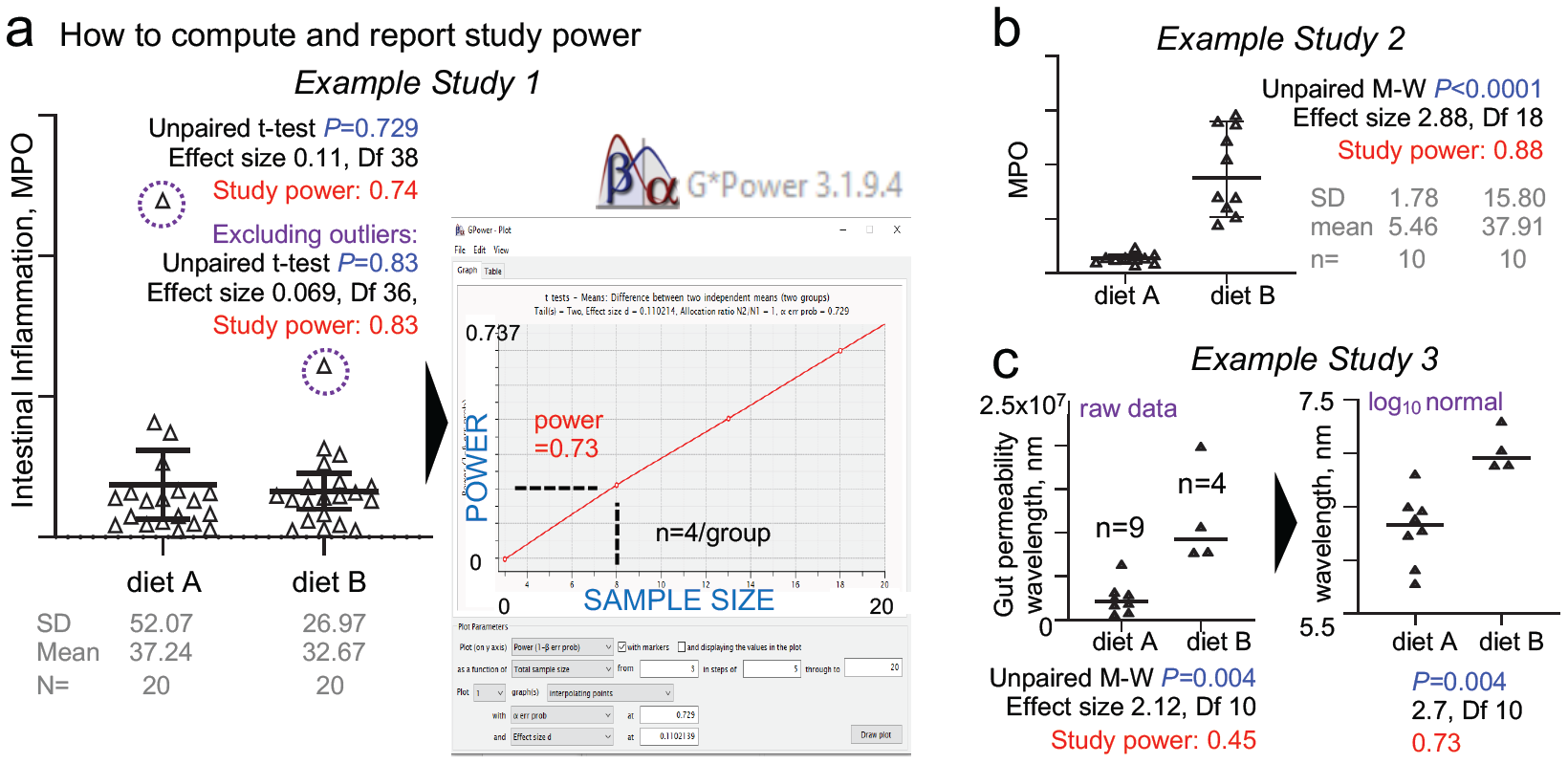
Graphical examples of rapid ‘study power’ calculations and reporting of individually-caged mouse data. **a**) Example of study power calculation & graphical reporting (post-hoc means after study completion, all datasets are real unpublished data). Intestinal inflammation in mice from two groups housed individually after pre-experimental cage microbiome normalization using IsPreFeH (fresh feces only; no bedding material). Post-test plot analysis (inset, software screenshot of power vs. sample size) shows that in this case, only 4 mice would be needed. Notice p-value and power increase after excluding outliers (dashed circles, N=19). **b**) Power analysis for two groups with different variance (diet A, narrow SD; diet B, wide SD). Fecal MPO test following a diet intervention illustrates that for this diet, a sample size of 10 is sufficient to achieve a well-powered study despite large variance in diet B. **c**) Example of importance of data normalization (e.g., from raw small changes in millions, 10^7^, to a log scale) in post hoc power analysis. Fluorescence intensity units in a test after intervention caused early mortality in diet B. Although the p-value does not change, normalized data (smooth edges of datasets) increases study power as it fulfills assumptions of t-tests normality. Since all mice were individually caged, the dataset quality and the early mortality are not due to, or are confounded by cage effects. Therefore, despite the small sample size (n=9 vs. 4), this is a well-powered study. The most recent version of open-source software G*Power can be downloaded from http://www.psycho.uni-duesseldorf.de/abteilungen/aap/gpower3/. See **Supplementary Figure 4** with step-by-step process to compute powers herein shown. Examples for power and sample size for studies with individually caged mice intended for ANOVA or regression analysis are available at https://stats.idre.ucla.edu/other/gpower/.

### Recommendation theme 6 on ‘Implementing statistical models to consider ICC in clustered data’

Depending on the experiment, we recognize that it is not always possible to single-house mice. Our review showed that scientists often analyze clustered observations using methods that mathematically function under the assumption of data independence (student T-, Mann-Whitney, One-/Two-way ANOVAs), without implementing statistics for intra-class (‘intra-cage’) correlated (ICC) cage-clustered data (Multivariable linear/logistic, Marginal, Generalized Estimating Equations, or Mixed Random/Fixed Regressions).^47,76,77^ The ICC describes how units in a cluster resemble one another, and can be interpreted as the fraction of the total variance due to variation between clusters.^47^ Housing multiple MxCg as homogeneous densities across study groups is logistically challenging using few cages. To expand the outreach of our multi-theme framework, and to support scientists with their analysis and publication of justifiable/clustered experiments, we **recommend** to ‘*<u>Use statistical methods designed for analyzing clustered data when multiple mice are housed in one cage, and when data points are obtained from mice over time, to **i)** properly assess treatment effects, **ii)** determine the intraclass correlation coefficient for each study, and then **iii)** to use that information to rapidly generate experiment-specific, customizable study power tables to aid in the assessment, re-/design (if more mice or cages are needed), and reporting of adequately powered studies.</u>’* The **goal** is to promote and facilitate the implementation of cage-clustered data analysis in mouse research by ***i)*** providing examples demonstrating the misleading effect when univariate methods are used for clustered-mice, and by ***ii)*** making our statistical code available to the public to gain familiarity with protocol principles of cluster-data statistical tools. Recommendation six is intended to serve as a technical guide supporting the framework, and therefore was not tested for implementability.

The statistical example we provide is based on data extracted (using ImageJ^78^ analysis) from a published dot plot figure in a reviewed study that exclusively reported cohousing 5 MxCg, and where authors compared two diets using 8 and 9 MxGr (2 TCgxGr; **Figure 9A**). The published p-value was 0.058, but to emphasize our message, we slightly/evenly adjusted the extrapolated data to achieve a univariate p<0.050. By simulating 5 possible cage-clustering scenarios, Figure 8 was designed to help visually understand the benefits of computing ICC and experiment-specific customizable power tables to determine whether more cages/group or mice/cage are needed to achieve study powers of ideally >0.8.

**Figure 9.**
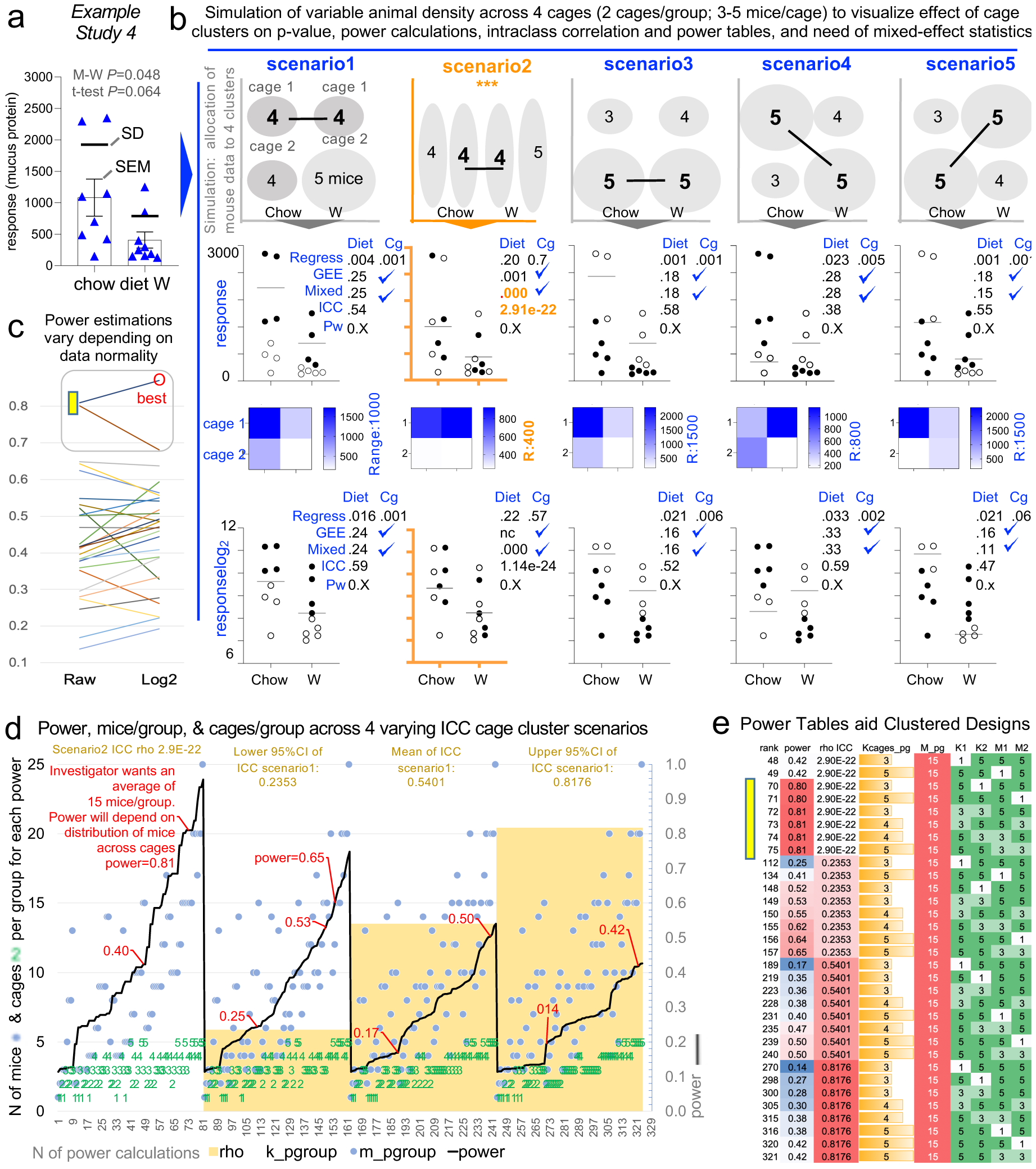
Analysis of cage-clustered data, intra-class correlation coefficients and power tables to facilitate study design by the number of cages/group and mice/cage. Five scenarios using a single dataset where mice housed as 5 mice/cage illustrate the effect of cage clustered data. Raw data extrapolated from one of the 172 reviewed studies. **a**) Extrapolated raw data (original published dot plot; p-value=0.059). Note that data from the diet ‘W’ group was not normally distributed (A-D p=0.005). **b**) Graphical representation of 5 scenarios considering different cage allocations of 2 cages/group. P<0.05 for the regression analysis (‘Regress’) indicates cage effect. Except for scenario 2 (with mice representing the entire data range spectrum for treatment outcome ‘response’; y-axis), note that all scenarios are subject to significant cage effect. ‘Diet’, treatment; ‘Cg’, Cage effect. **c**) Paired line plot depicting power estimates for the same data, without transformation (left) and after log2 transformation (right). Note the best power estimation for raw data may have a marked influence on the power estimation based on log2. We advise log2 transformed data for this dataset. **d**) Line plot depicting power calculations for three ICCs, as a function of ICC, (histogram). Power estimations depend on ICC, simulation determined the n MxCg and TCgxGr needed to achieve a study power, which changes depending on degree of cage clustering (i.e., ICC, intra-class correlation coefficients). **e**) Power table illustrates the number and distribution of mice using a clustered design to achieve study power.

When using clustered-data methods, we showed that only one of the five scenarios yielded a significant diet treatment effect (*i.e.*, scenario 2, where all cages were unbiased, having mice with high and low response values, something unlikely to occur naturally in clustered settings, **Figure 9B**). Data proves that artificial heterogeneity due to mouse caging and unsupervised ‘cage-effects’ lead to poor reproducibility (80% of cases would misleadingly show that the test diet induces an effect on the mouse response). Graphically, we show that the variability of ICC (computed after running the mixed-effect models) depends on the hypothetical mouse allocation to cages, which in turn influences the post-hoc estimations of study power (**Figure 9C-D**).

As a final practical product in this manuscript, we provide the statistical scheme/code in the GitHub repository (https://github.com/axr503/cagecluster_powercode) to implement this streamlined analysis and compute comprehensive power tables based on the ICC derived for each simulation to help scientists determine the best mice-to-cage combinations to match resources (**Figure 9E**). A ‘quick reference’ of actionable steps for all six themes is in **Supplementary Table 8**. To expand our implementability strategy for continuous assessment by the international scientific community the survey is available online (https://forms.gle/LxPCydbySddcndZ7A).

## DISCUSSION

This study and proposed framework were motivated by the identification of a wide heterogeneity in published methods relevant to diet, microbiome, and the pathogenesis of inflammatory bowel diseases and digestive health in humans, where mice models are critical to study diseases biology, translational interventions, or to inform clinical trials for humans. The actionable framework described however applies to any field of modern mouse research. Although it is impossible to develop a single consensus statement on practices pertaining to experimental supplies (*e.g.*, bedding type, water, facilities) to accommodate every scientific community/goal, our proposed implementability statistics indicate that the 6-theme actionable framework described could be widely adopted to reduce the deleterious impact of these emerging concepts on artificial heterogeneity. This framework especially designed around reducing animal density, cage dirtiness, and cage microbiome bias, stresses the need of statistical methods for power and cluster data.

Herein, we confirmed that research methodology continues to vary in published literature, and as documented by a survey of academicians, such variability may be attributed to well-ingrained heterogeneous perceptions among scientists concerning how animal husbandry impacts the mouse microbiome. Animal density and the cost dilemma of how many cages are used to test hypotheses in experiments were deemed amenable for improvement. Because adjustments to facility settings are not easy to standardize, we propose that the most experimentally effective strategy to improve study power/reproducibility in the literature is to implement a lower number of mice per cage. From our analyses, we provide recommendation themes to minimize cage-clustering effects and implement clustered data analysis methods as a means to reduce artificial heterogeneity.

Although adding more cages to a study increases handling costs, studying *‘less mice per cage is more’* is a pro-statistically powerful, comparably effective practice. The use of cost as a rationale for conducting cage-clustered experiments needs conscientious consideration, since housing costs are just a fraction of the research funds required for tests. Perhaps, institutions could provide discounts to investigators for the cost of housing when conducting experiments, because fewer MxCg requires less cage changes and experiments are often short-term. Logistically, since fewer MxCg may be an option limited by space in certain facilities, well-powered and well-analyzed cage-cluster studies is desirable.

**In conclusion**, we confirmed that research methodology continues to vary in published literature and as documented by a survey of academicians. Analyses indicate that the reporting of post-hoc study power calculations, in the context of the proposed framework, could be objectively used to guide and monitor the research power and reproducibility across mouse microbiome research at large.

## MATERIALS and METHODS

### Study overview

As an overall methodological strategy to confirm and quantify the extent to which animal husbandry variability has been, and continues to be, present in mouse and microbiome research, we *<u>first</u>* conducted a quantitative verification of animal husbandry variability in academia ***i)*** by screening the recent published peer-reviewed literature (2018-2019) to infer the historic prevalence of prevailing practices that could have influenced research and ***ii)*** by conducting a survey of academicians across relevant professional organizations to determine the present status on beliefs and knowledge on husbandry practices. *<u>Then</u>*, we ranked the practices based on relevance to influence microbiome research, as perceived by respondents, to prioritize/make six recommendations. *<u>Lastly</u>*, to document the validity of such recommendations, we conducted a targeted literature search to cite examples enabling the analysis of such suggestions in future consensus efforts. Using a Delphi-based consensus strategy, these suggestions were graded for quality to compute heterogeneity and probability statistics for implementability by investigators. See Figure 1 for illustrated study overview.

### Quantification of husbandry methods heterogeneity

As a test topic, we chose to use ‘dietary studies in mice’ as PubMed search terms to screen (scoping review) original peer-review studies for animal husbandry practices as of May 3^rd^, 2019, published literature (see references of identified studies in **Supplementary Materials**) To interrogate and quantify perceptions and opinions among academicians on animal husbandry practices that influence microbiome data variability, a one-time online IRB-approved survey with 11 multiple-choice questions was administered, via recruitment email, to eligible participants through membership list servers of the following: ***i)*** faculty of 17 NIH National Institute Diabetes and Digestive and Kidney Diseases (NIDDK) Silvio O’Conte Digestive Diseases Research Core Centers (‘DDRCC’), which provide research support to local and national institutions, ***ii)*** registrants of the 2018 Cleveland International Digestive Education and Science (IDEAS) Symposium hosted by the Cleveland DDRCC, Case Western Reserve University (CWRU), ***iii)*** registrants of the Taconic Biosciences Webinar titled ‘Cyclical Bias and Variability in Microbiome Research’, ***iv)*** members of the American Association of Laboratory Animal Science (‘AALAS’), and ***v)*** members of ‘GNOTOBIOTIC’ ListServ, forum of the National Gnotobiotics Association.

### Six evidence-based recommendations graded for future implementability

To provide evidence-based suggestions and to support the development of a large-scale consensus report that can be implemented and beneficial to research, we used a ranking of the survey-derived husbandry practices to prioritize the husbandry topics deemed influential in mouse microbiome by respondents. Using Google PubMed and keywords contained in the survey question/topic (*e.g.*, mouse, water), five coauthors cataloged relevant peer-reviewed scientific articles on each topic (targeted review). The information gathered, as tables, was used as assessment tools by 14 individuals to grade a table with 5-recommendations drafted by the lead and senior authors in this study. Collectively, the individuals comprised professional experiences across five research institutions; CWRU, The Scripps Research Institute, Kyorin University, South Dakota State University, The Ohio State University, University of Chicago, and Cornell University. To determine if the 5-recommendations could be implemented as a framework, individuals were asked to provide suggestions, new recommendations, and to grade (1, low; 10, highest) each item for sentence clarity, potential impact, and recommendability to others (**Supplementary Materials**). These ‘implementability grades’ numerically illustrate the potential for variance and adoption of the recommendations by others in mouse research.

### Ethical considerations

All research was approved by the Case Western Reserve University Institutional Review Board (STUDY20180138).

### Statistics

For computation purposes, animal/cage density data extracted from the scoping review were used to create a secondary index. Specifically, the number of animals per group (group size, MxGr) and the number of mice housed per cage (animal density, MxCg) were used to compute a semi-descriptive index metric of ‘cage cluster effect’ on each study: ‘estimated number of cages per experimental group’ (*i.e.*, total n of cage clusters per group, TCgxGr = MxGr divided by MxCg). If a range was provided for animal density (*e.g.*, 1-5), estimations were computed using the median value within the range, as well as the minimum and maximum values. Average of estimated center values were used for analysis and graphical summaries. For Figure 9, study selection was based on the use of 5 mice/cage, and that study results were published as dot plots (allowing us to infer the raw data for our analysis) in the manuscript. Descriptive statistics for parametric data were employed if assumptions were fulfilled (*e.g.*, 1-way ANOVA). Non-fulfilled assumptions were addressed with nonparametric methods (*e.g.*, Kruskal-Wallis). As needed, 95% confidence intervals are reported to account for sample size (*e.g.*, MxCg; surveyed participants) and for external validity context. Significance was held at p<0.05. Analysis, study powers, and graphics were conducted with R, STATA, Python 3.0 Anaconda, GraphPad and G*Power.^74^ G*Power is an open-source power specialized software for various family of tests; calculations only require p-value (alpha), sample size, and mean±SD to compute effect size. Excel was used to create a cage handling frequency and cost spreadsheet calculator.

## Abbreviations

AALAS: American Association of Laboratory Animal Science;
DDRCC: Digestive Diseases Research Core Center;
GF: germ-free;
ICC: intra-class correlation coefficient;
IsPreFeH: Inter-subject Pre-experimental Fecal Microbiota homogenization;
MxCg: mice per cage;
MxGr: mice per group;
TCgxGr: total cages per group.

## ACKNOWLEDGMENTS

Authors want to express their gratitude to Mrs. Colleen Karlo at Case Western Reserve University for her guidance during survey preparation and institutional guidance to ensure compliance and protection of the human subjects that participated in the survey. The authors are indebted to all the survey participants for their time and suggestions. Jonathan Craven for his administrative support and the IDEAS symposium participants. Special thanks go to Drs. Jung-Fu Chang, DVM, PhD, Population Medicine, Cornell University; Joy Scaria, PhD, Animal Diseases Research and Diagnostic Laboratory, South Dakota State University; Craig L. Franklin, DVM, PhD, Mutant Mouse Resource and Research Center, Department of Veterinary Pathobiology, University of Missouri-Columbia; Eugene B. Chang, MD, PhD, Department of Medicine and Microbiome Center, University of Chicago, for enlightening discussions, comments and scientific suggestions in various phases of this study which ultimately influenced the directions and focus of the present report.

## AUTHORS CONTRIBUTIONS

AB and ARP envisioned and planned this study, conducted the survey and data analysis, interpreted the data, and wrote initial manuscript draft. AL DK and GL conducted literature searches under the direction of AB and ARP, read the manuscript, and provided critical comments for evidence based-factors. All authors were involved in providing comments and discussing the six recommendations drafted by the lead and senior authors. All authors reviewed and commended on the final recommendations. ARP and SI implemented the grading strategy of recommendations. ARP conducted statistical analysis and power analysis, outlined excel calculator, and wrote statistical scripts. ELM provided suggestions and verified statistical scripts. FC and external MS, JM and BRT were major contributors in interpretation, rationale for outreach of study and contributed with suggestions during manuscript preparation. All authors approved the final manuscript.

